# Vessels hiding in plain sight: quantifying brain vascular morphology in anatomical MR images using deep learning

**DOI:** 10.1101/2025.05.06.652518

**Authors:** Asa Gilmore, Anita Esi Eshun, Yue Wu, Aaron Lee, Ariel Rokem

## Abstract

Non-invasive assessment of brain blood vessels with magnetic resonance (MR) imaging provides important information about brain health and aging. Time-of-flight MR angiography (TOF-MRA) in particular is commonly used to assess the morphology of blood vessels, but acquisition of MRA is time-consuming and is not as commonly employed in research studies or in the clinic as the more standard T1- or T2-weighted MR contrasts (T1w/T2w). To enable quantification of brain blood vessel morphology in T1w/T2w images, we trained a neural network model, anat2vessels, on a dataset with paired MR/MRA. The model provides accurate segmentations as assessed in cross-validation on ground truth images, particularly in cases where T2w images are used. In addition, correlation between features that are extracted from model-based vessel segmentations and from ground truth account for as much as 78% of the variance in these features. We further evaluated the model in another dataset that does not include MRA and found that `anat2vessels`-based vessel morphology features contain information about aging that is not captured by cortical thickness features that are routinely extracted from T1w/T2w images. Moreover, we found that vessel morphology features are associated with individual variability in blood pressure and cognitive abilities. Taken together these results suggest that `anat2vessels` could be fruitfully applied to a range of existing and new datasets to assess the role of brain blood vessels in aging and brain health. The methods are provided as open-source software in https://github.com/nrdg/anat2vessels/.

## 1. Introduction

Brain aging is a multifaceted process that includes many different aspects that can be measured with magnetic resonance imaging (MRI). In the white matter, a wide array of senescence effects includes reduction in the overall volume, which can be imaged with structural T1- and T2-weighted MRI (T1w/T2w) [1], as well as specific changes to the microstructure, composition, and geometry of white matter connections, which can be imaged with diffusion-weighted and quantitative MRI [2]. The gray matter also changes its volume and composition, which can be quantified via estimates of cortical thickness and quantitative MRI measurements [3]. Many of the cellular changes to neurons and glia that affect these measurements occur in tandem with changes to brain blood vessels [4]. Brain blood vessels change their physical properties with aging due to a variety of mechanisms, including atherosclerosis and arterioscelrosis, with measurable changes in their gross morphology. Vascular risk factors that affect these changes, such as diabetes [5], hypertension[6, 7] and atrial fibrillation [8] are also associated with dementia risk in aging. Therefore, non-invasive assessments of brain blood vessel morphology is an important tool in studying brain aging. Time of flight magnetic resonance angiography (TOF MRA) offers a view of large brain blood vessels and their morphology, which is useful in clinical assessment of brain infarcts and other gross dysfunctions of the vascular system, such as Moyamoya disease and cerebral vasculitis [9]. In addition, these measurements provide information about aging-related changes to blood vessel morphology [10]. Features of brain blood vessel morphology from TOF-MRA also provide information about systemic cardiovascular phenotypes associated with hypertension [11].

Studying the role of changes to vascular morphology in cognitive aging and neu-rodegenerative diseases is presently limited, because many studies that pertain to brain aging do not collect MRA images. Fortunately, information about brain vessel morphology is also visible in other MRI contrasts, such as T1w and T2w. However, information about blood vessel morphology is not routinely extracted and quantified from these measurements in standard neuroimaging processing pipelines, such as Freesurfer [12], because of the lack of image analysis tools that extract these features. Here, we developed a novel tool – the `anat2vessels` neural network – that segments and quantifies brain blood vessel morphology in routinely-collected T1w/T2w MR images. This model was trained on a dataset (IXI) in which MRA was collected in tandem with T1w and T2w images, allowing us to learn in a data-driven manner about the relationships between these images. We evaluated the model both in the IXI dataset, in which it was trained (using held-out test sets), as well as in a different dataset (CamCan) in which no MRA is available. This evaluation demonstrated that the model is accurate, can capture substantial variance in the quantitative features of blood vessel morphology, as well as substantial variance in aging that is not captured in routine evaluation of T1w/T2w images. In the absence of an MRA, this method also captures substantial variance in individual characteristics of high relevance to aging research: cognition and blood-pressure.

## 2 Results

### 2.1 Assessing model accuracy in the model development dataset

In the IXI dataset, `anat2vessels` can accurately produce vessel segmentations. An example (Figure 1) from held-out test data demonstrates that this accuracy corresponds to a close spatial match between ground truth (green) and model-generated vessel segmentations.

**Fig. 1:**
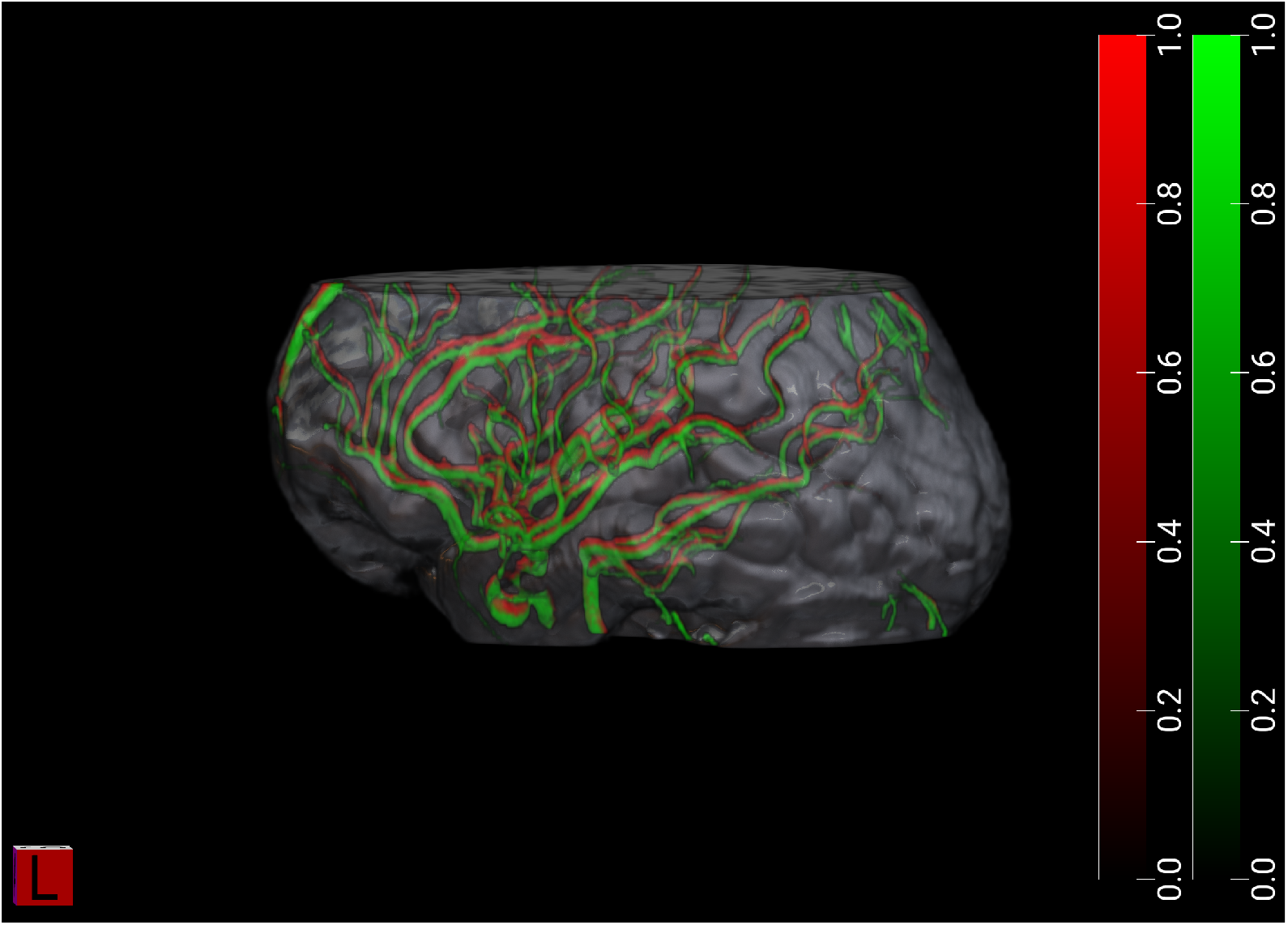
Example from one IXI test-set subject (IXI265), demonstrating accurate brain blood vessel segmentation. Ground truth (COSTA) in green, and model segmentation in red. This example corresponds to a Dice coefficient of 0.28.

In 5-fold cross-validation, model accuracy is similar between T2w and T1w + T2w input data, but is substantially lower for only T1w input data (Figure 2). Dice scores are, for T1w + T2w: 0.55 ± 0.14 (SD), for T1w only: 0.37 ± 0.14, and for T2w only: 0.52 ± 0.15 (SD). Nevertheless, Dice is significantly higher for the T1w+T2w model both in comparison to the T1w model (*t*_426_=26.6, *p <*0.001) and in comparison to the T2w model (*t*_426_=1.97, *p <*0.05). Model accuracy is not related to age: correlations between Dice coefficients and age are: *R*^2^ = 0.01 (T1w only), *R*^2^ = 0.02 (T2w only) and *R*^2^ = 0.01 (T1w+T2w). Because the model trained with T1w and T2w provides the most accurate results, all subsequent analyses are based on the outputs of the T1w+T2w model.

**Fig. 2:**
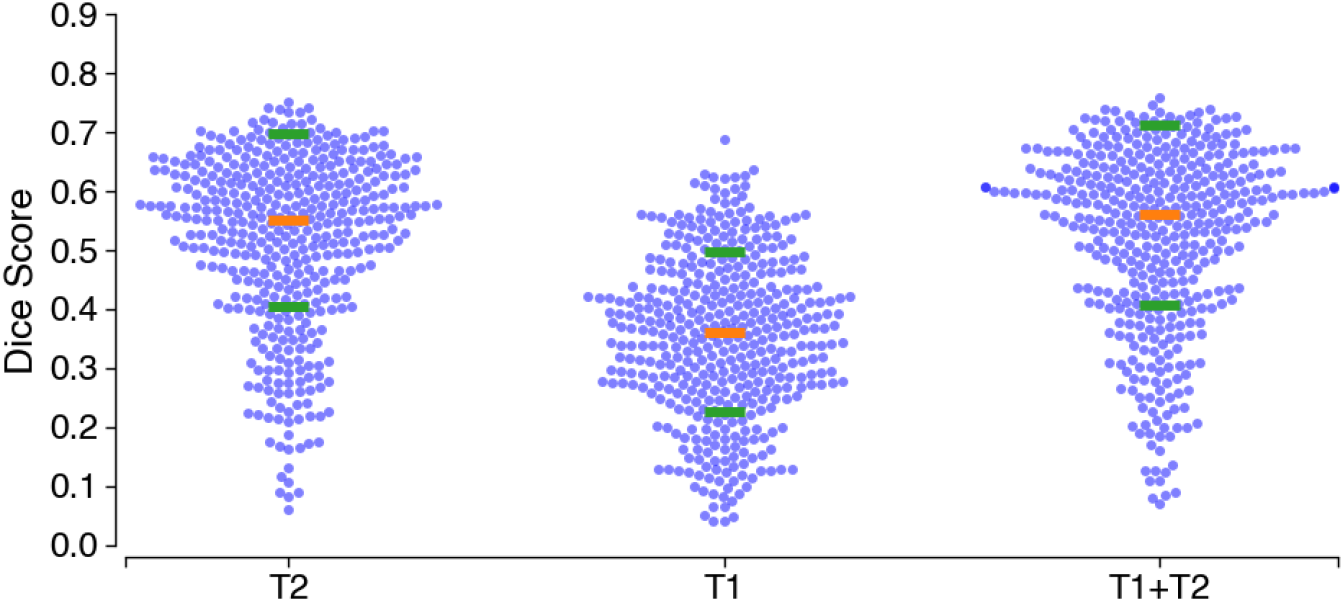
Dice score for each participant in the IXI data set in a 5-fold cross-validation procedure. Orange horizontal bars indicate the median and green bars are ± standard deviation

While mean Dice accuracy is not very high, features extracted from the ground-truth vessel masks do have very high correlations with morphological features extracted from the model-generated vessel masks (Figure 3). For example, model mean radius (Figure 3E) accounts for 78% of the variance in ground-truth mean radius across individuals in the test-set. Total volume (Figure 3B) from the model accounts for 67% of the variance in the volume in ground-truth extracted from MRA images.

**Fig. 3:**
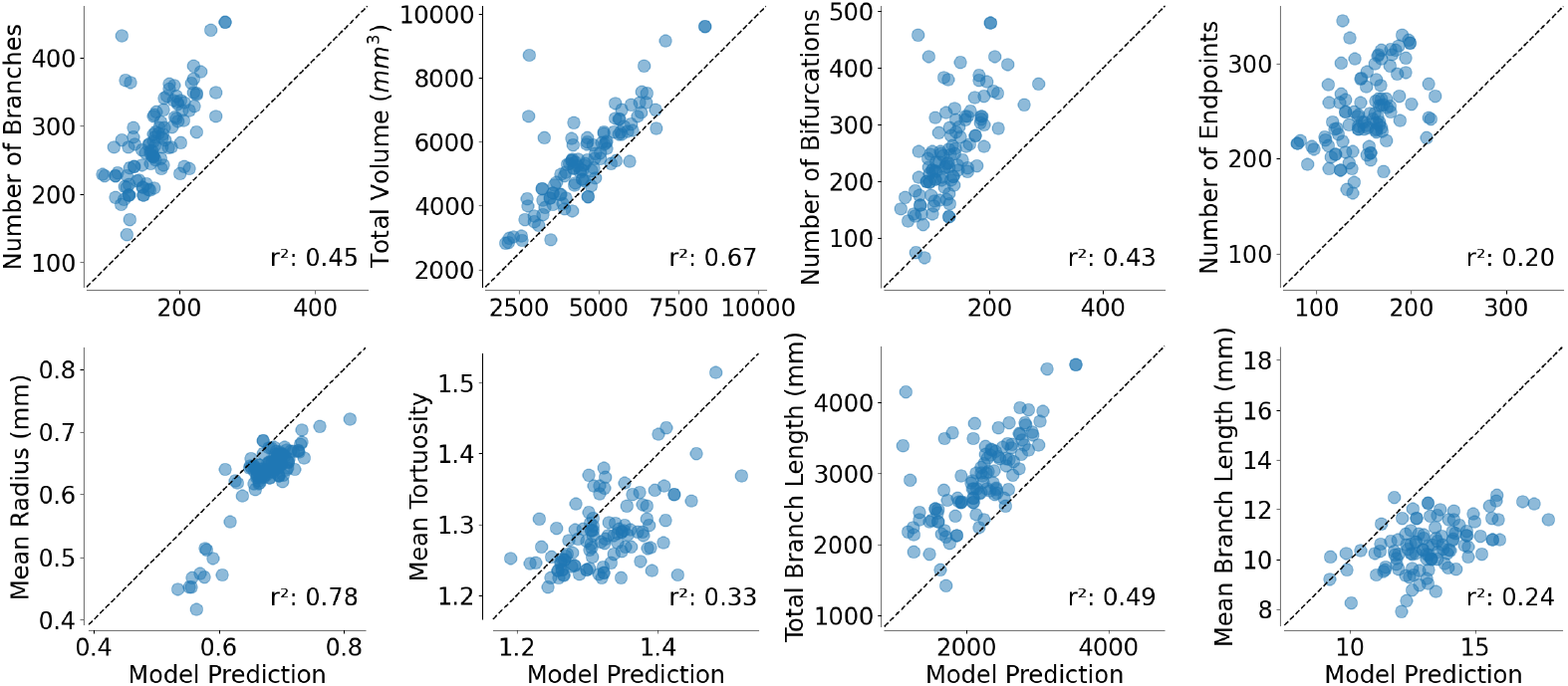
Vessel features extracted from T1w+T2w model predictions (abscissa) compared to those extracted from COSTA segmented MRA masks (ordinate). Here, *r*^2^ indicates variance explained, quantified as squared Pearson correlation coeffcients.

Consistent with previous research, when quantitative morphological vessel features are combined using a linear model, they account for substantial variance in age. In the test set, a linear model that uses ground truth vessel features has an adjusted *R*^2^=0.35. The same model, fit to `anat2vessels`-derived features has an only slightly lower adjusted *R*^2^=0.31. Moreover, the residuals of age models fit to ground-truth and `anat2vessels`-derived features are highly correlated across the individuals in the test set (Pearson *r*^2^ = 0.67, *p <*0.001). Furthermore, there is no correlation between the squared-residuals of the model fit to `anat2vessels`-derived features and the accuracy of the `anat2vessels` network segmentation, quantified as the Dice coefficient (Pearson *r*^2^ = 0.01, p=0.2). This suggests that the limiting factor in sensitivity to age-related changes is not due to the same factors that limit Dice accuracy.

The accuracy of the age model based on `anat2vessels`-derived features was also verified in a 10-fold cross-validation procedure. In each iteration, the model was fit to 90% of the data and predicted the left out 10%. This procedure was repeated 10 times, such that every participant had their age predicted with a model that was fit to a training dataset in which their data was not included, so one *R*^2^ value can be calculated that includes the full sample. A similar proportion of variance explained was found with this procedure (Figure 4; cross-validated coefficient of determination *R*^2^ = 0.3). Here as well, `anat2vessels` segmentation accuracy, quantified as Dice coefficients (color of data points in Figure 4) does not correlated with the error in modeling age using these features (Pearson *r*^2^ = 0.0005, p=0.6).

**Fig. 4:**
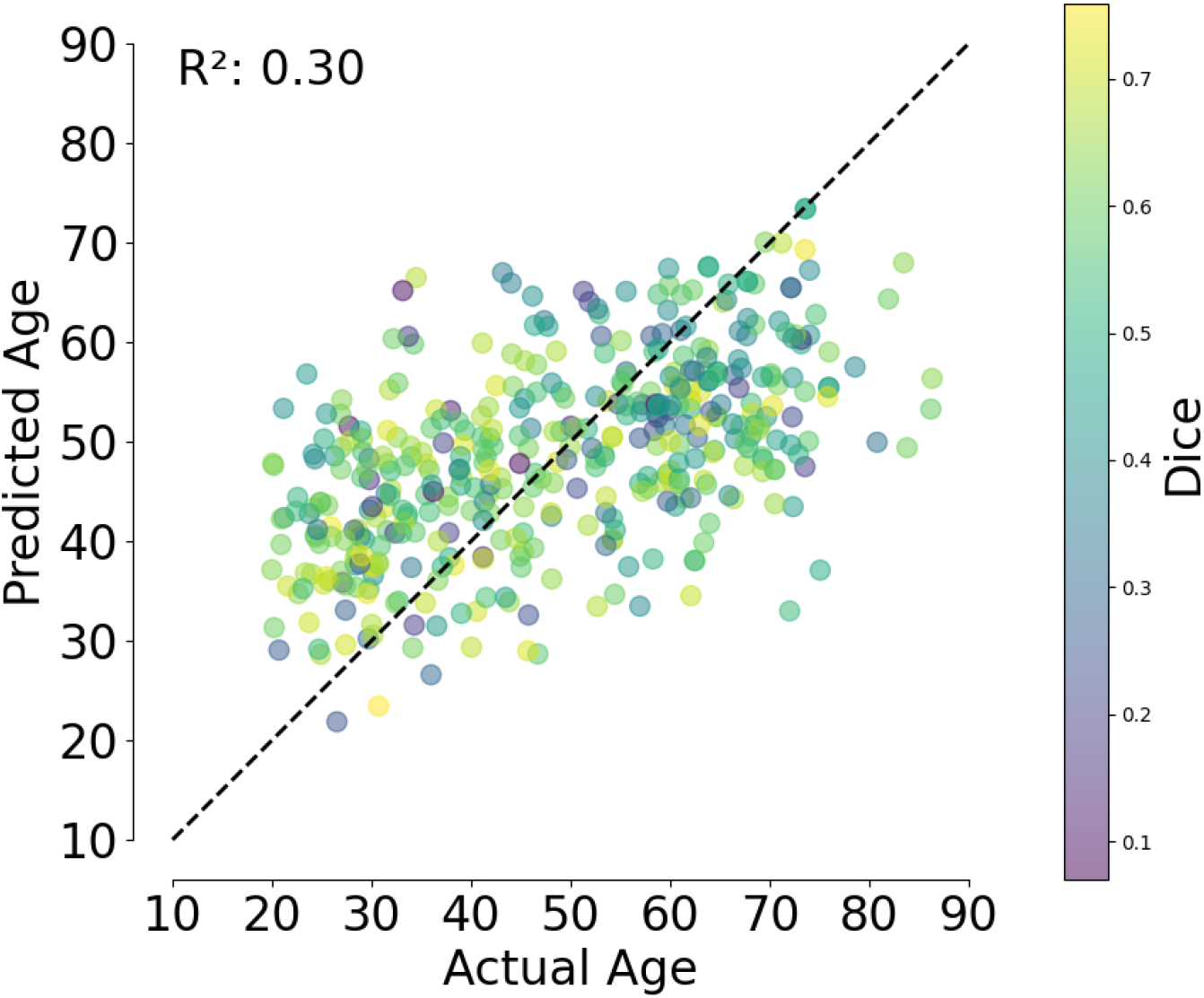
Age prediction on IXI dataset using 10-fold cross validation with an OLS model, colored by Dice score.

### 2.2 Evaluating the model in an independent dataset

To assess the generalization of these results to data acquired with different scanners, and different acquisition parameters, we used a separate dataset: the CamCan life-span dataset. In CamCan, TOF-MRA were not collected, so it is not possible to assess the accuracy of vessel mask segmentation directly. Instead, we assessed the contribution of `anat2vessels`-based features to models of invdividual variability in several characteristics. First, we asked whether blood vessel features derived with `anat2vessels` can improve models of brain aging. A model of age as a linear combination of vessel features, similar to the one fit to the test set in the IXI dataset has an adjusted *R*^2^ of 0.39, confirming that these features are informative on their own.

As a baseline to evaluate whether `anat2vessels`-based features provide information that was not previously available from anatomical images, we also fit a model to cortical thickness features computed with Freesurfer. Consistent with a large body of research on brain aging, this model accounts for a substantial portion of variance in aging, with an adjusted *R*^2^ of 0.6. However, the variance explained by these two models does not fully overlap. We can demonstrate this by fitting a model that contains both cortical and vessel features. This model has a substantially higher adjusted *R*^2^ of 0.69. An analysis of variance between the cortex-only model and the cortex-and-vessels model finds that the models differ significantly in terms of their explained variance, even when accounting for the difference in the number of parameters (*F*_11,575_=13.5, *p <*0.001). We also repeated this comparison using a 10-fold cross-validation procedure. Here, *R*^2^ for the vessels-only model was 0.41, the cortex-only model had a cross-validated *R*^2^ of 0.53, and the cortex-and-vessels model had an *R*^2^ of 0.62 (Figure 5).

**Fig. 5:**
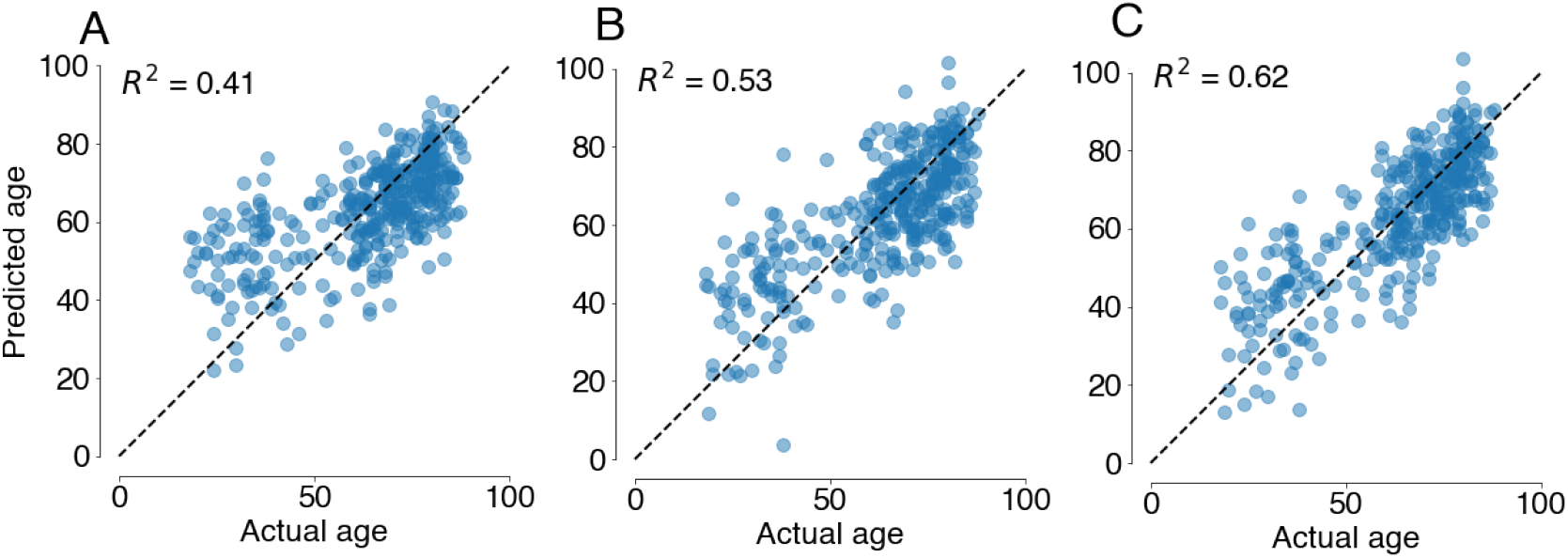
Age prediction on CamCan dataset using 10-fold cross validation for models that include only cortical thickness features (**A**), only vessel features(**B**), and both (**C**). Here, variance explained, *R*^2^ is quantified as the coefficient of determination.

Finally, we tested whether vessel features could be used to explain individual differences in participant attributes related to cardiovascular and cognitive aging. Again, for each dependent variable: diastolic and systolic blood pressure and performance in the MMSE, we compared a model with cortex-only features (and with age and sex as covariates) to a model with cortex-and-vessel features. We found that vessel features significantly improved the model for systolic blood pressure (*F*_11,485_=2.37, *p <*0.01), but not for MMSE or diastolic blood pressure (both *p >*0.05). However, when the data was split into older (above 60 years old) and younger (60 and under) participants, we found that the addition of vessel features improved the model of MMSE in older individuals (*F*_11,186_=1.93, *p <*0.05) and the addition of vessel features improved modeling of both diastolic (*F*_11,264_=1.91, *p <*0.05) and systolic (*F*_11,264_=3.13, *p <*0.001) blood pressure in younger individuals. Furthermore, we identified individuals with hypertension in the CamCan dataset regardless of their age, defined as diastolic blood pressure above 90 mmHg or systolic blood pressure above 140 mmHg (Figure 6). A logistic regression model that includes both cortex and vessel features classifies these individuals significantly better (Figure 6, *F*_11,650_=3.1, *p <*0.001).

**Fig. 6:**
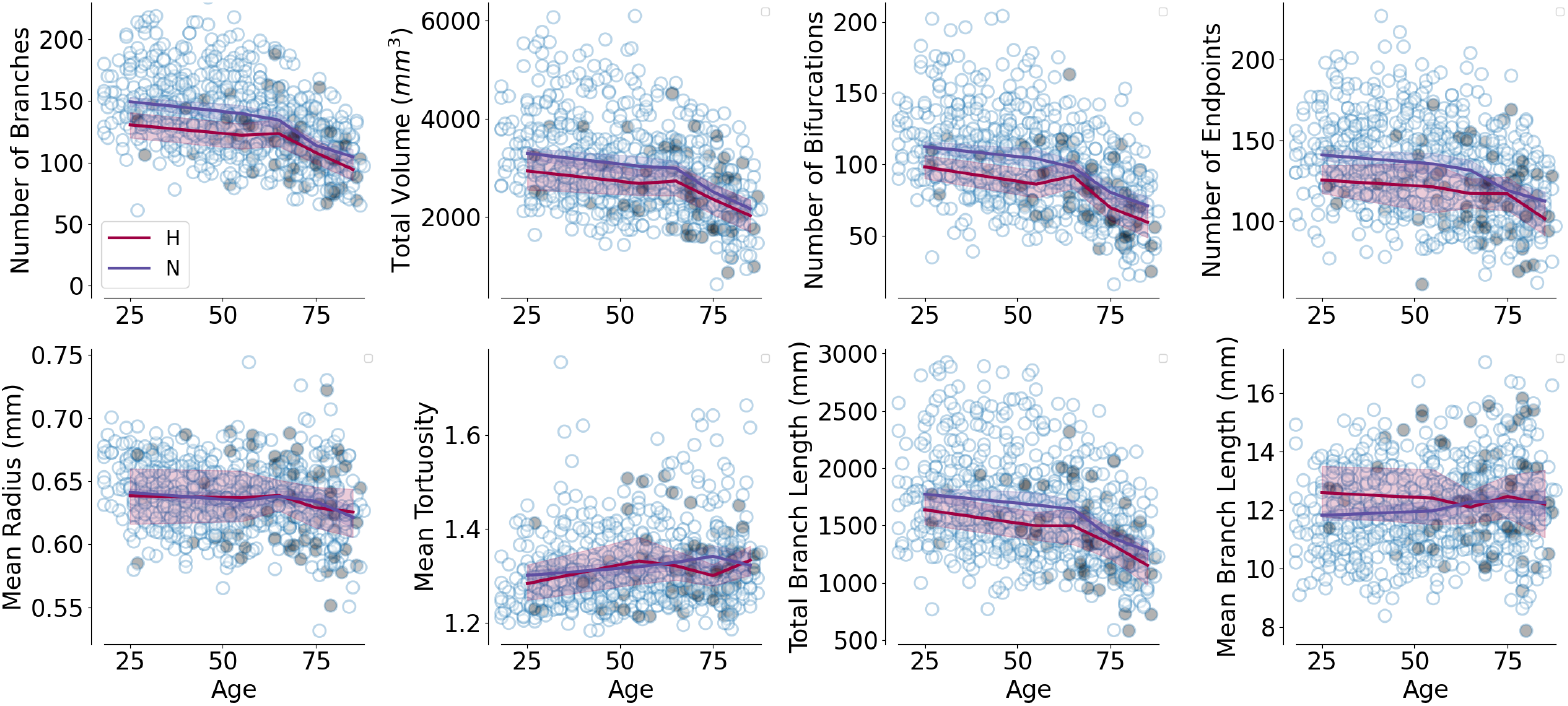
Age dependence of vessel features in the CamCan dataset. Individuals with hypertension are identified as filled circles. Lines indicate age-bin averaged features for hypertensive (“H”) and noromotensive (“N”) individuals, with bootstrapped 95% CI (age bins: *<* 50, 50 − 60, 60 − 70, 70 − 80, *>* 80)

**Fig. 7:**
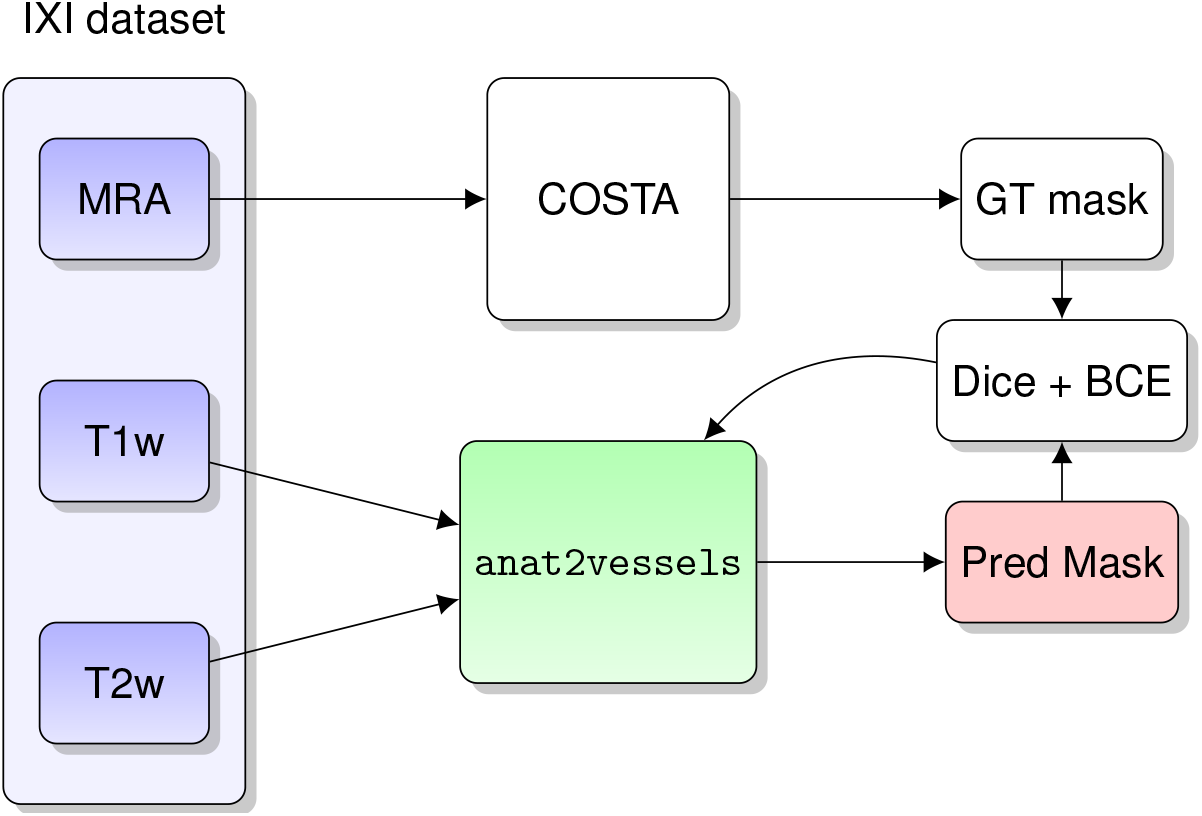
The `anat2vessels` training pipeline.

**Fig. 8:**
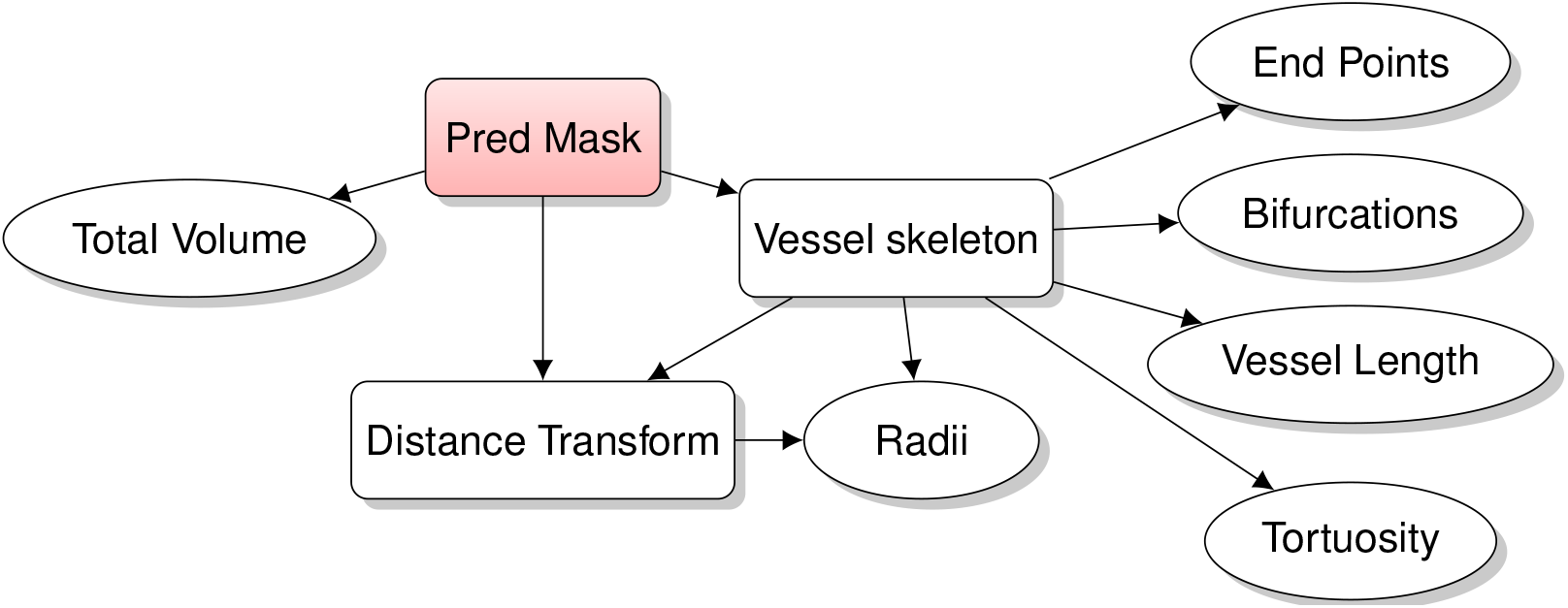
Feature Extraction pipeline.

**Fig. 9:**
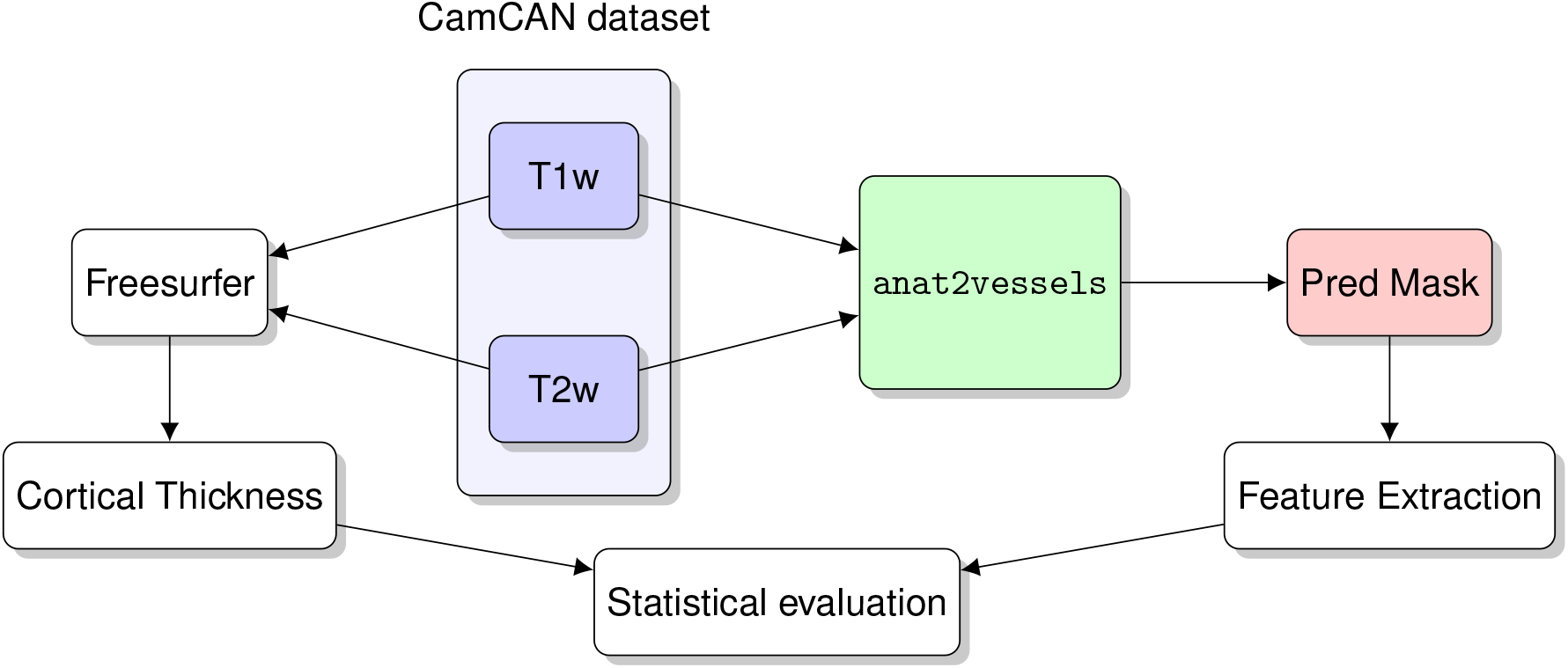
Validation pipeline.

## 3 Discussion

There is an increasing interest in understanding the role that brain vasculature plays in cognitive aging and brain health. However, dedicated vascular imaging can be prohibitively expensive and onerous and in many already existing datasets was not included, thus limiting the ability to ask questions about brain blood vascular morphology. The present work advances studies that incorporate this information by providing a method to extract morphological features of brain blood vessels from commonly acquired anatomical imaging.

To be broadly useful, a method for blood vessel morphology quantification needs to fulfill several criteria: (1) It needs to be accurate. We found that the model that we developed has good accuracy with respect to gold standard segmentation of blood vessels in MRA. Moreover, while features extracted from the segmentation are biased relative to ground truth – for example, the total volume of `anat2vessels` segmentation is almost always smaller than ground truth (Figure 3) – they are highly correlated with features extracted from ground-truth vessel segmentations, suggesting that they are capturing meaningful variability between individuals. (2) It needs to relate to known causes of brain blood vessel change. For example, we demonstrate that vessel features in the IXI dataset account for substantial variance in subject age, known to be related to changes in brain blood vessels [4]. Furthermore, the residuals of a model based on `anat2vessels` features are highly correlated with the residuals of a model based on features calculated from ground-truth vessel segmentation, providing further evidence that the `anat2vessels` segmentations capture true biological variability. (3) It needs to generalize to datasets that were acquired on other MRI instruments. Computer vision models based on deep learning are notoriously sensitive to domain shift [13]. Therefore, we evaluated the model in a second dataset in which no TOF MRA was collected. While this dataset does not provide a gold-standard benchmark, it provides several silver-standard evaluation targets: evaluating the known relationship between features of blood vessel morphology and age, as well as assessments of relationships between synthetic blood vessel morphology features and either blood pressure or cognitive abilities.

We found that: (1) vessel morphology features from the `anat2vessels` model demonstrate the expected link with age; (2) this link with age is not fully explained by cortical thickness features derived from the anatomical data; and (3) model features are linked to both cognitive abilities (particularly within older individuals) and to blood pressure (particularly within younger individuals). These results dovetail with previous findings that show linkages between cardiovascular health and cognitive aging [6–8].

### 3.1 Limitations and future work

One limitation of this work is that features derived from `anat2vessels` are biased estimates of vessel features. For example, the volume of the vessels is uniformly underestimated. This means that, while they are highly informative about inter-individual variability, the features derived with `anat2vessels` are not accurate as absolute quantitative assessments of brain blood vessel features.

One of the challenges of the training data that we used is the limited field of view of the MRA data. This means that the model generalizes well to other scanners/datasets, but not outside of the circumscribed anatomical field of view that was used for training (exemplified in Figure 1). A limitation of CamCan as an evaluation dataset is the limited range of MMSE scores: subjects were excluded from MRI measurements if their MMSE was lower than 24. This means that we are examining a limited range of cognitive deficits. In addition, there is a limited number of individuals with hypertension. Using the standard definition of hypertension as diastolic blood pressure higher than 90 or systolic blood pressure higher than 140, we find only 75 individuals of the 650 in this dataset that meet this criterion (Figure 6). Future work will focus on datasets where MRI measurements were conducted in tandem with substantial cognitive decline and/or cardiovascular conditions. This will allow the method to uncover as yet undetected linkages between brain vasculature and cognitive decline in aging.

Future work could be undertaken to increase the anatomical specificity of the conclusions presented here. For example, there is some evidence that certain parts of the vasculature display more age effects than others [14]. Therefore, extensions of this work will focus on delineating vessel properties that are identified into different vascular territories and/or functional groups.

### 3 2Conclusions

This technical report introduces a novel tool for quantification of blood vessel tissue properties in neuroanatomical images, which provides information about individual differences in blood vessel morphology features. The tool that we developed explains variance in age and in other characteristics linked to brain blood supply. Taken together, these evaluations demonstrate that the method has potential for high utility across a range of different datasets pertaining to brain aging, vascular health and the intersection between them. Code and model weights are openly available in https://github.com/nrdg/anat2vessels.

## 4 Methods

### 4.1 Data

We used two datasets in our analysis. The first dataset was used to train the `anat2vessels` neural network model and the second dataset was used to evaluate `anat2vessels` performance in assessment of brain aging and modeling individual differences. Both datasets are anonymized, publicly available datasets, and the University of Washington IRB does not consider access to anonymized publicly available neuroimaging datasets to be human subjects research.

#### 4.1.1 Training dataset: IXI

The openly available IXI dataset (https://brain-development.org/ixi-dataset/) was used to train our model on T1w, T2w and blood vessel masks extracted from matching MRA scans. The dataset contains 582 subjects, of which 568 have T1w, T2w, and MRA imaging present (ages: 19-86, 277 male) with images acquired in 3 different scanners. For a breakdown of acquisition parameters for each scanner see Table 1.

**Table 1:** MRI Acquisition Parameters for IXI Dataset.

| Scanner | Modality | TR (s) | TE (ms) | In-plane Resolution (mm) | Slice Thickness (mm) |
| --- | --- | --- | --- | --- | --- |
| IXI: Phillips Intera 3T (185 subjects) | T1w | 9.6 | 4.6 | $0.94 \times 0.94$ | 1.2 |
| | T2w | 5.7 | 100 | $0.94 \times 0.94$ | 1.2 |
| | MRA | 16.7 | 5.75 | $0.47 \times 0.47$ | 0.8 |
| IXI: Phillips Gyroscan Intera 1.5T (322 subjects) | T1w | 9.8 | 4.6 | $0.94 \times 0.94$ | 1.2 |
| | T2w | 8.2 | 100 | $0.94 \times 0.94$ | 1.2 |
| | MRA | 20.0 | 80 | $0.47 \times 0.47$ | 0.8 |
| IXI: GE 1.5T (75 subjects) | T1w | N/A | N/A | $0.94 \times 0.94$ | 1.2 |
| | T2w | N/A | N/A | $0.94 \times 0.94$ | 1.2 |
| | MRA | N/A | N/A | $0.26 \times 0.26$ | 0.8 |
| CamCan: Siemens 3T TIM Trio (653 subjects) | T1w | 2.25 | 2.99 | $1.0 \times 1.0$ | 1.0 |
| | T2w | 2.8 | 408 | $1.0 \times 1.0$ | 1.0 |

#### 4.1.2 Evalutation dataset: CamCan

The evaluation dataset was accessed through the Cambridge Centre for Ageing and Neuroscience (Cam-CAN) archive [15]. Here, 653 subjects (ages 18-88, 322 male) were available, for which only T1w and T2w MRI are available. See Table 1 for parameters of the acqusition.

### 4.2 Models

The IXI dataset was used to train a model to produce binary segmentations of vessels from T1w/T2w inputs. For every subject in this dataset, blood vessel segmentation masks were obtained using the COSTA cerebrovascular segmentation deep learning network, which was trained on a multi-site, multi-vendor TOF MRA dataset [16]. This algorithm produces binary masks of the vascular tree for each input MRA image at an identical resolution to the input. We chose the COSTA model after comparing with an expert-segmented dataset [17], as we found that it produced more reliable and accurate results.

We trained the nnU-Net ResEnc L model [18, 19] on paired T1w/T2w/T1w+T2w and blood vessel segmentations derived from the MRA images. T1w and T2w images were resampled to the MRA resolution with linear interpolation using nibabel [20].

For training and evaluation, we randomly split the IXI dataset into train (N=427) and test (N=106) samples, with an even split of each scanner in each set. Following standard nnUNet procedure, 5-fold cross-validation was conducted within the training set, stratified by scanner.

The model was trained with an initial learning rate of 0.01, and linear decay to zero over 1,000 epochs. The batch size was 35, and the patch size was 64× 320 ×320. Each model for the 5-fold cross validation was trained on an NVIDIA H100 NVL 94gb GPU. Training took aproximately 29 hours per fold. For precise details on the training procedure, see the nnUNet documentation [18], as well as the “plans.json” file in the accompanying software repository.

### 4.3 Feature extraction

To produce interpretable and anatomically based predictions of age and disease from the vessel segmentation’s produced by `anat2vessels` we extract a number of features from the masks. To do so we first produce a “skeleton” of the mask using the Lee thinning algorithm [21]. This is used to define the center-line of each blood vessel, as well as to compute the number of bifurcation points and endpoints in the vessel map. We can also compute an estimate of the radius at each point by computing a distance transform from the center-line to the edge at each point on the centerline. This centerline is then broken up by Bifurcations into a set of individual vessels. We then measure the total length of each vessel as well as the length between the endpoints of the vessel. We use these measurements to calculate tortuosity for each vessel as: 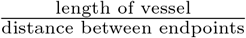. Finally, we calculate the total volume of a vessel mask by summing the number of voxels in the mask. All measurements take into account the actual size of the vessels. A Python script for extracting these features is available in the repository for this paper.

### 4.4 Validation experiments

After training, we used a separate dataset, the CamCAN dataset, to evaluate `anat2vessels` performance. Because the CamCAN dataset does not contain MRA scans, vessel features were extracted from the `anat2vessels`-generated vessel segmentations. To run inference on the CamCAN dataset we first run synthstrip to remove the skull [22], and then perform a rigid registration with a reference image in the IXI dataset. We then used the feature extraction pipeline above to extract vessel features from `anat2vessels`. We used Freesurfer version 7 run on T1w and T2w data to also extract cortical thickness in cortical regions of the Desikan Killiany parcellation.

#### 4.4.1 Statistical evaluation

Several models were fit to the data, to evaluate the use of `anat2vessels`-derived vessel features. Sensitivity to age was assessed with the following models (using Patsy formula notation [23]):

> *Age* ∼ regional cortical thickness
>
> *Age* ∼ vessel features
>
> *Age* ∼ regional cortical thickness + vessel features

which where subsequently used to predict age in a 10-fold cross-validation procedure. For each of diastolic and systolic blood pressure, and MMSE, the following models were fit:

> *DV* ∼ age + C(sex) + regional cortical thickness
>
> *DV* ∼ age + C(sex) + vessel features
>
> *DV* ∼ age + C(sex) + regional cortical thickness + vessel features

where “DV” is the dependent variable, *C*(*sex*) indicates that this variable is included as a categorical variable. In addition, participants were classified as having hypertension if they had diastolic blood pressure above 90 mmHg or systolic blood pressure above 140 mmHg. Further logisitic regression models were fit to discriminate individuals with and without hypertension:

> hypertension ∼ logistic(age + C(sex) + regional cortical thickness)
>
> hypertension ∼ logistic(age + C(sex) + vessel features)
>
> hypertension ∼ logistic(age + C(sex) + regional cortical thickness + vessel features)

All statistical models were fit using the Python Statsmodels library [24] and comparisons between nested models were conducted using analysis of variance functionality implemented in this library. Cross-validation also used sample splitting functionality implemented in Scikit Learn [25].

## 5 Availability

All code for this paper has been made available on GitHub. A csv file containing the subjects used for training and testing has also been added to the repository. The IXI dataset can be found at (https://brain-development.org/ixi-dataset/). The CamCAN dataset is not a public dataset, but those interested can apply for access at (https://camcan-archive.mrc-cbu.cam.ac.uk/dataaccess/).

## 6 Acknowledgements

This work was funded by National Institutes of Health grants MH121868, MH121867, EB027585, R01AG060942, and U19AG066567, as well as by National Science Foundation grant 1934292 and by Research to Prevent Blindness.

Preliminary testing for this work was completed on Hyak, UW’s high performance computing cluster. This resource was funded by the UW student technology fee.

Data collection and sharing for this project was provided by the Cambridge Centre for Ageing and Neuroscience (CamCAN). CamCAN funding was provided by the UK Biotechnology and Biological Sciences Research Council (grant number BB/H008217/1), together with support from the UK Medical Research Council and University of Cambridge, UK.

Thanks to Kendrick Kay and Aviv Mezer for helpful discussions of this work.

